# Tenascin-C activation of lung fibroblasts in a 3D synthetic lung extracellular matrix mimic

**DOI:** 10.1101/2023.02.24.529926

**Authors:** Aritra Nath Kundu, Carey E. Dougan, Samar Mahmoud, Alara Kilic, Alexi Panagiotou, Ninette Irakoze, Nathan Richbourg, Shelly R. Peyton

## Abstract

The lung extracellular matrix (ECM) maintains the structural integrity of the tissue and regulates the phenotype and functions of resident fibroblasts. Lung-metastatic breast cancer alters these cell-ECM interactions, promoting fibroblast activation. There is a need for bio-instructive ECM models that contain the ECM composition and biomechanics of the lung to study these cell-matrix interactions *in vitro*. Here, we developed a synthetic, bioactive hydrogel that mimics the native lung modulus, and includes a representative distribution of the most abundant ECM peptide motifs responsible for integrin binding and matrix metalloproteinase (MMP)-mediated degradation in the lung, which promotes quiescence of human lung fibroblasts (HLFs). Stimulation with transforming growth factor β1 (TGF-β1), metastatic breast cancer conditioned media (CM), or tenascin-C activated these hydrogel-encapsulated HLFs in a manner reflective of their native *in vivo* responses. We propose this lung hydrogel platform as a tunable, synthetic approach to study the independent and combinatorial effects of ECM in regulating fibroblast quiescence and activation.

## 1. Introduction

The lung ECM plays an important role in regulating the phenotype and function of the resident cells, and in maintaining healthy tissue homeostasis. Fibroblasts represent the primary cell type in the lung ECM, and they are responsible for the deposition of most lung ECM proteins (predominantly collagens I and III, elastin, and fibronectin)^1^, and their turnover via collagenase, stromelysin, gelatinase, and plasminogen activator. Conversely, fibroblast phenotype, migration, proliferation, and differentiation are regulated by biochemical and biophysical cues from the surrounding lung ECM. This bi-directional fibroblast-ECM interaction is integral to the regulation of connective tissue morphogenesis and dynamics during homeostasis and wound healing^2–4^. For example, during wound healing, the ECM protein fibrin enables the migration of fibroblasts to wounds and directs production of collagen to initiate repair^5^. Integrin β_1_ expressed by lung fibroblasts drives fibroblast migration via focal adhesion kinase (FAK), and subsequent collagen I deposition and wound repair ^6–8^. Perturbations in this cell-ECM interaction may lead to ECM remodeling and stiffening, characteristic of fibrosis in lung tissue.

In healthy lung tissue, fibroblasts are quiescent, but become activated by changes in lung ECM stiffness^9,10^. Fibroblasts cultured on high-tension collagen matrices assume a “synthetic” phenotype, associated with activation and proliferation^2,11–13^. Fibroblast activation in response to ECM stiffening is associated with a myofibroblastic phenotype, characterized by prominent stress fibers rich in α-smooth muscle actin (α-SMA)^14–19^. The fate of the ECM is intricately linked with fibroblast activation. As one example, during breast cancer metastasis, interactions between the ECM and fibroblasts establish a “feed-forward” amplification loop in which myofibroblasts produce matrix, and the matrix activates signaling pathways that support fibroblast activation and survival^20,21^. Breast cancer metastasis promotes production of transforming growth factor β1 (TGF-β1), a pro-fibrotic cytokine, which promotes myofibroblast differentiation and survival in conjunction with the matrix-mediated signals^22–24^. Several *in vitro* platforms have been developed to study fibroblast phenotype and activation in stiff substrates, mimicking fibrotic environments^25^, largely decellularized tissue scaffolds^26^ and collagen gels^2,5,27,28^. We sought to create a synthetic *in vitro* lung ECM to study fibroblast phenotype in the context of lung tropic breast cancer metastasis. Synthetic hydrogels provide a tremendous opportunity to design a lung-specific scaffold with nuanced control over ECM components, which has not been done to study quiescence and activation of lung fibroblasts.

To do this, we first characterized the human lung ECM via mechanical indentation, mass spectrometry, and Protein Atlas histology^29^ to incorporate the appropriate modulus and most prevalent ECM protein composition responsible for integrin-mediated adhesion and matrix metalloprotease (MMP)-mediated degradation in the lung. We synthesized peptides to represent these proteins and combined them with a modulus-tunable poly(ethylene glycol) (PEG) network to create a carefully designed, yet very simple synthetic lung ECM hydrogel. With this hydrogel, we retained primary human lung fibroblast quiescence *in vitro* and, achieved precise control over fibroblast activation in response to TGF-b1 and metastatic breast cancer. and tenascin-C.

## 2. Results

### 2.1 Design of synthetic lung ECM mimic

We used a previously published approach to identify the key ECM properties of lung tissue that could be represented in a synthetic, PEG-based hydrogel (Figure 1)^30,31^. To identify the ECM proteins in real lung tissue, we referenced published information on proteins expressed in the lung from the Protein Atlas (citation here), as well as our own quantitative mass spectrometry on human lung samples. We performed liquid chromatography-mass spectrometry (LC-MS/MS) on ECM-enriched healthy lung tissue from six different donors (Figure 1a). This full list of matrisome proteins was then narrowed to those that mediate cell attachment via integrins (21) or are susceptible to cleavage by matrix metalloproteinases (MMPs, 27). Then, we identified the specific peptide sequences within these ECM proteins that are either responsible for high-affinity binding to integrins (Figure 1b)^2^ or are highly susceptible to cleavage by MMPs (Figure 1c). The integrin-binding peptides were synthesized with a single cysteine for covalent immobilization within the hydrogel (Figure 1b)^32–38^, and the MMP-degradable peptides were synthesized with cysteines on each end to act as crosslinkers (Figure 1c). To determine the relative molar concentrations for each peptide included in the synthetic lung hydrogel, we devised a scoring system inspired from scoring systems previously described in our bone marrow and brain mimicking hydrogels (Figure 1b, c)^30,31^.

**Figure 1.**
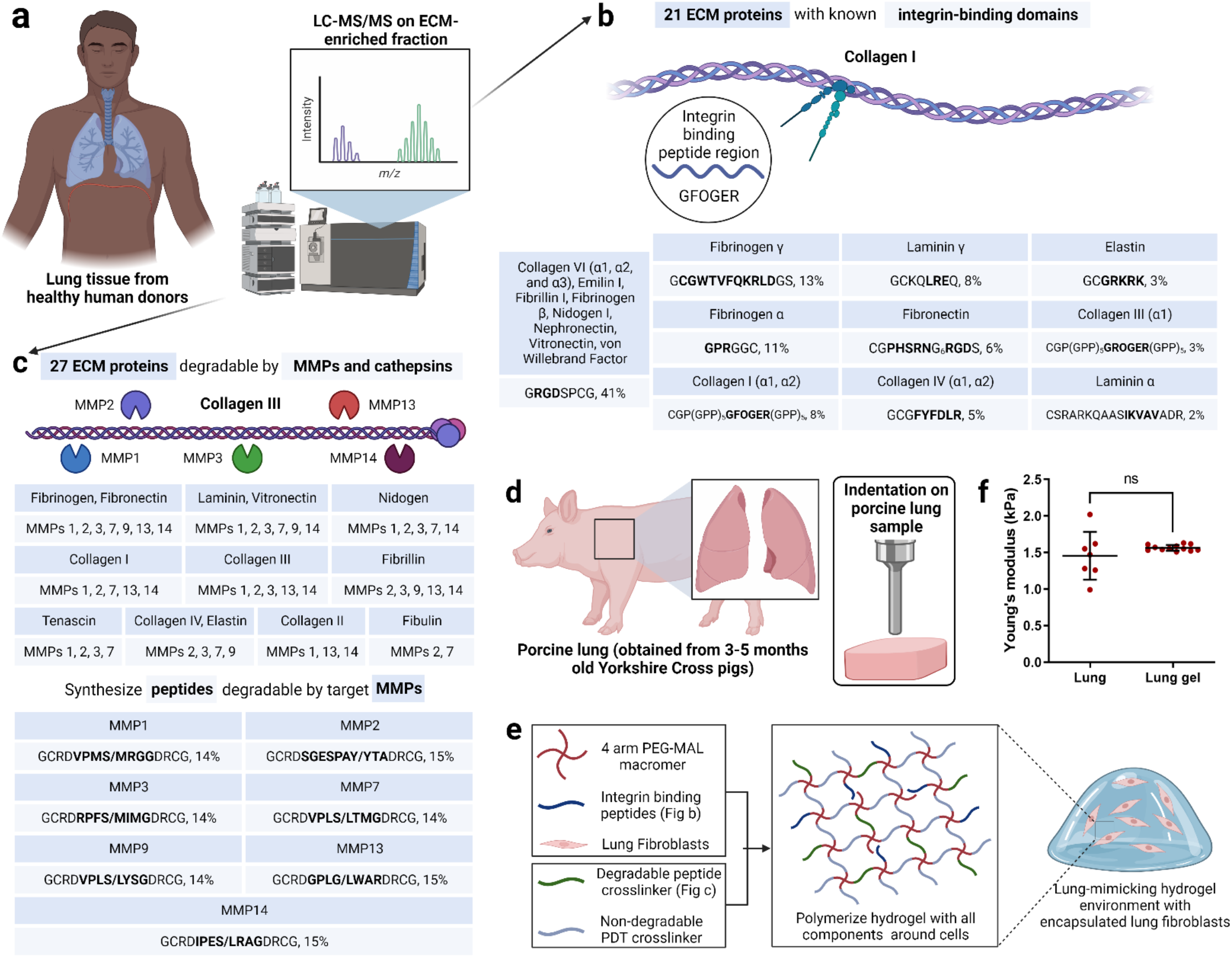
Design of synthetic 3D lung ECM hydrogel. a) Schematic of characterization of lung ECM using LC-MS/MS on ECM-enriched lung tissue samples from healthy human donors. b) Illustration of integrin-mediated binding to collagen I and the list of identified integrin-binding peptide domains with their relative quantities corresponding to ECM proteins. c) Illustration of MMP-mediated degradation of collagen III and the list of MMPs responsible for ECM protein degradation along with the list of identified MMP-degradable peptide domains with their relative quantities corresponding to the different MMPs. d) Schematic of mechanical characterization of porcine lung using indentation method. e) Schematic of lung ECM hydrogel preparation with encapsulated lung fibroblasts. f) Young’s modulus of lung hydrogel measured using indentation method and its comparison with the lung modulus using the same method.

In our previous work, we quantified the modulus of porcine lungs via indentation, shear rheology, uniaxial tension, and cavitation rheology^39^. We used this lung modulus (1.4±0.4 kPa) to guide the crosslinking density of our PEG hydrogel. We compared the compressive modulus of porcine lungs and the different formulations of our PEG hydrogel (Figure 1d, Supplementary Figure S1a). Upon careful evaluation of multiple hydrogel parameters that can be used to manipulate hydrogel modulus, a 10 wt%, 4-arm, 10kDa PEG hydrogel, with a modulus of 1.5±0.4 kPa was selected to match the reported modulus of porcine lung tissue (Supplementary Figure S1b).

One benefit of synthetic hydrogels is that their moduli can be independently tuned from the concentration of bioactive peptides included. To ensure this was the case for our hydrogel, which includes 17 different peptides, we individually incorporated the peptides into the hydrogel and tested their effects on modulus. Replacing MMP-degradable peptide crosslinkers with 1.5 kDa PEG-dithiols up to 25 mol% did not alter the hydrogel modulus (Supplementary Figure S1c). Integrin-binding peptides could be incorporated up to a 2 mM total concentration without compromising the bulk modulus of the hydrogel (Supplementary Figure S1d). In sum, our final lung-mimicking hydrogel formulation from 4-arm, 10 kDa PEG-maleimide at 10 wt% in solution, functionalized with 2 mM integrin-binding peptides and crosslinked at molar equivalence to available maleimide groups with a 75:25 (molar ratio) mixture of PEG dithiols and MMP-degradable peptides, with a modulus of 1.5±0.1 kPa (Figure 1e, f).

### 2.2 Functional validation of lung ECM peptides

We adapted a cell attachment assay to validate the attachment of human lung fibroblasts (HLFs) to peptides at the molar concentrations used in the lung hydrogel (Supplementary Figure S2a)^40^. We observed a significant increase in the cell surface area on glass coverslips functionalized with integrin-binding peptides (Figure 2a, Supplementary Figure S2b). The extent of cell spreading was dependent on the concentration of the peptide with the highest cell attachment observed on the complete integrin-binding peptide cocktail. We also performed a competitive cell adhesion assay to further validate binding to integrin peptides^40^. This involved seeding HLFs pre-treated with individual peptides (soluble in the cell culture medium) onto surfaces that were covalently coated with the full integrin-binding peptide cocktail (Supplementary Figure S3a) and measuring cell size over two hours. When cells were pre-treated with soluble peptides, we observed a significant decrease in cell area on the coverslip, which we surmise is from the peptides in solution competing for integrin receptors on the cell membrane (Supplementary Figure S3b).

**Figure 2.**
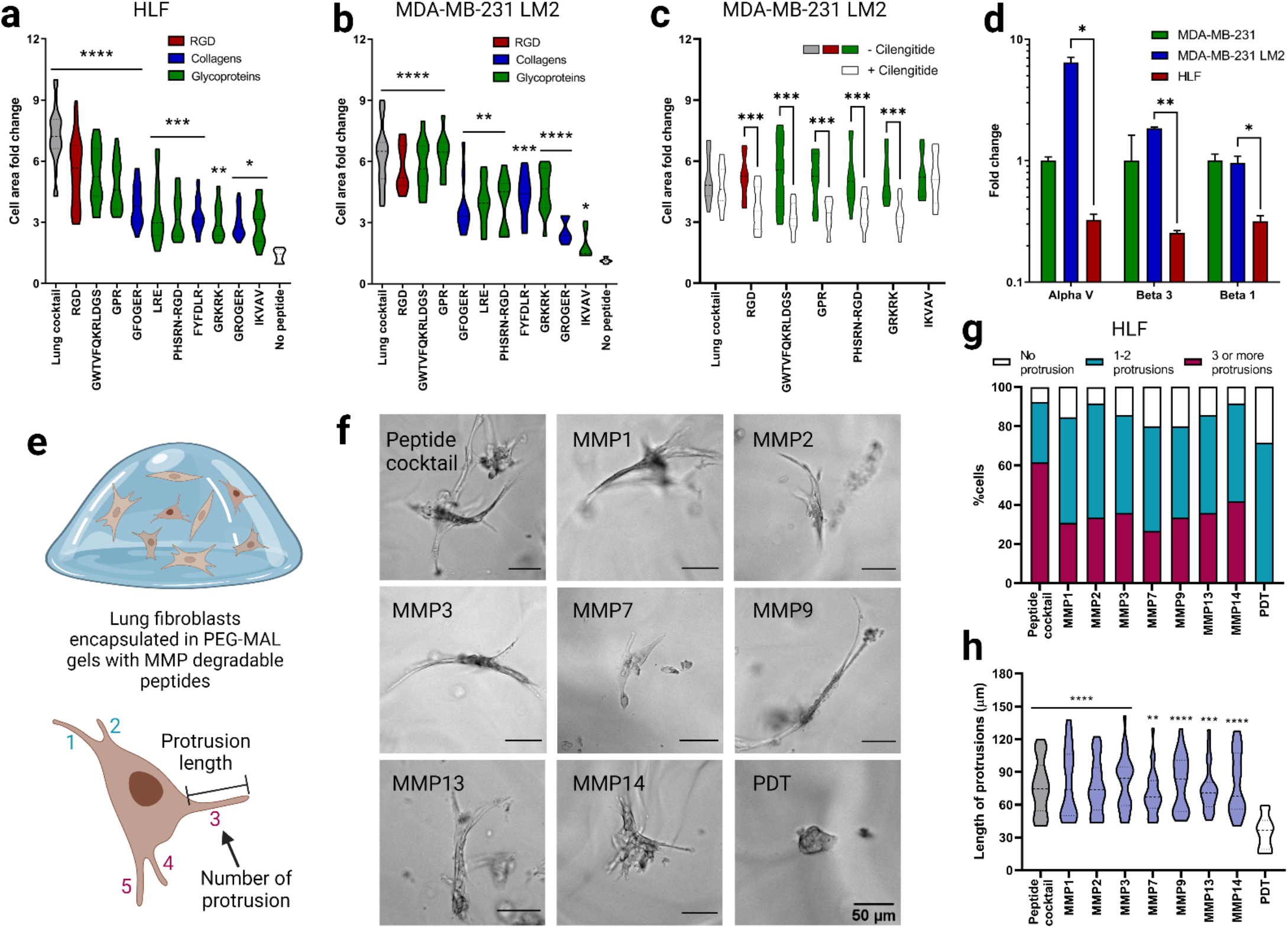
Functional validation of lung integrin-binding and MMP-degradable peptides. a) Cell area fold change 2 hours after seeding human lung fibroblasts (HLFs) onto glass coverslips functionalized with intergrin-binding peptides relative to a negative control (glass coverslip treated with silane but no peptides). b) Cell area fold change for lung tropic MDA-MB-231 breast cancer cells in comparison to negative control. c) Cell area fold change on a subset of the integrin-binding peptides in the presence of cilengitide, reported in comparison to positive control (no cilengitide). d) Fold change in gene expression, measured via qRT-PCR, for αv, β3 and β1 integrins in lung tropic MDA-MB-231 breast cancer cells and HLFs, each gene is normalized to the parental MDA-MB-231 breast cancer cells. e) Illustration of HLFs in 3D hydrogels and measurement of cell protrusions quantification. f) Representive images of HLFs after 6 days of encapsulation. g) Quantification of HLF populations with different number of protrusions for individual MMP degradable peptides and lung MMP degradable peptide cocktail. h) HLF protrusion length for individual MMP degradable peptides and lung MMP degradable peptide cocktail. All data are mean + s.d. Statistical analyses were performed using Prism (GraphPad). Data in (a), (b), (c), (d), (g) and (h) were analyzed using a one-way analysis of variance followed by a Dunnett’s multiple comparison test with 95% confidence interval. *, **, ***, and **** indicate P < 0.05, P < 0.01, P < 0.001, and P < 0.0001.

We then further examined the affinity of breast cancer cells including lung tropic triple-negative breast cancer MDA-MB-231 LM2 cells, parental MDA-MB-231 cells, and HER2+ BT474 cells toward the lung integrin-binding peptides. While MDA-MB-231 is a highly aggressive and invasive breast cancer cell line, BT474 is known to be a non-invasive breast cancer cell line with very low metastatic potential to the lung. A significant increase in cell area was observed for MDA-MB-231 LM2 cells, indicating increased cell attachment to the integrin-binding peptides compared to the negative control coverslip without any peptides (Figure 2b, Supplementary Figure S2c). Although peptide concentration-dependent cell attachment to the integrin-binding peptides was observed, MDA-MB-231 LM2 cells attached significantly more to fibrinogen γ (GWTVFQKRLDGS) and elastin (GRKRK) peptides than other peptides with similar concentration, such as collagen I (GFOGER) and collagen III (GROGER) peptides, respectively. A similar trend was observed for the parental MDA-MB-231 cells (Supplementary Figure S2d, e). However, non-lung tropic BT474 cells did not show significant attachment to the lung peptides (Supplementary Figure S2f, g).

HLFs primarily express integrins α_2_β_1_, α_3_β_1_, and α_v_β_3_ and other α_v_ and β_1_ containing heterodimers^6,8,12^. MDA-MB-231 cells, on the other hand, overexpress integrin α_v_β_3_. Several ECM proteins including fibrinogen α, fibrinogen γ, fibronectin, and elastin have a high affinity to integrin α_v_β_3_. To tease out the integrin α_v_β_3_ mediated cell attachment to peptides, we incubated the MDA-MB-231 LM2 cells with cilengitide, a cyclic RGD peptide that inhibits integrin α_v_β_341_. Cilengitide treatment significantly affected MDA-MB-231 LM2 cell attachment to RGD, elastin (GRKRK), fibrinogen α (GPR), fibrinogen γ (GWTVFQKRLDGS), and fibronectin (PHSRN-RGD) compared to untreated cells (Supplementary Figure S4). To further examine the α_v_β_3_ integrin-mediated cell attachment to these individual peptides, glass coverslips were functionalized with individual cilengitide-sensitive peptides (RGD, GRKRK, GPR, GWTVFQKRLDGS, PHSRN-RGD), along with the complete integrin-binding peptide cocktail, and laminin α peptide (LRE) as a negative control. Cells pre-incubated with cilengitide showed reduced cell attachment to the individual cilengitide-sensitive peptides (RGD, GRKRK, GPR, GWTVFQKRLDGS, PHSRN-RGD) compared to the non-incubated cells (Figure 2c). Cilengitide incubation did not affect the cell attachment to the laminin α peptide (LRE) (Figure S4b). RT-PCR results revealed increased expression of α_v_, β_3_ and β_1_ integrins in MDA-MB-231 LM2 cells compared to HLFs (Figure 2d), supporting the cell-specific adhesion results to the lung-specific integrin-binding peptides chosen for the synthetic lung hydrogel.

Incorporated MMP-degradable peptides were validated using a cell degradation assay with HLFs. HLFs are characterized by their elongated morphology and long protrusions, and we expected that gels without MMP-degradable peptides would yield rounded HLF morphologies and/or small protrustions. HLFs were encapsulated in hydrogels containing integrin-binding and MMP-degradable peptide cocktail at the same molar ratio of thiol to maleimide (Figure 2e). HLFs were able to extend processes in hydrogels that incorporated both integrin-binding and MMP-degradable peptides and remained viable in the lung hydrogel for 7 days (Figure 2f, g). Further quantification of HLF process length from the cell center, showed that HLFs in degradable hydrogels were able to extend their processes over 7 days, unlike HLFs cultured in hydrogels without degradable peptides (Figure 2h). To validate that each of the MMP-degradable peptides was susceptible to degradation by HLFs, cells were encapsulated in hydrogels containing individual MMP-degradable peptide crosslinkers or the complete MMP-degradable peptide cocktail. When degradable peptides were present, HLFs were able to branch further into the surrounding network (Figure 2f, h). Collectively, these results demonstrate that the HLFs can degrade lung hydrogels containing integrin-binding and MMP-degradable peptide cocktails while maintaining healthy cell morphology for more than a week.

### 2.3 HLF activation via TGF-β1

As a control, we first quantified the phenotype of lung fibroblasts on 2D first, we cultured HLFs on tissue culture polystyrene (TCPS) in serum-free fibroblast basal medium (FBM). After 6 days, we observed low α-SMA and FAP staining, indicating low activation of the HLFs (Figure 3a, b, c, Supplementary Figure S5a, c, d). Low expression of Ki67 further validated the quiescence of the HLFs on 2D (Figure 3f, Supplementary Figure S5f). Non-activated, quiescent fibroblasts assume a bipolar spindle-shaped morphology, which we also observed in HLFs cultured on TCPS (Supplementary Figure S5a). We observed the same quiescent phenotype in the 3D lung hydrogels with similar spindle-shaped morphology (Figure 3a-f).

**Figure 3.**
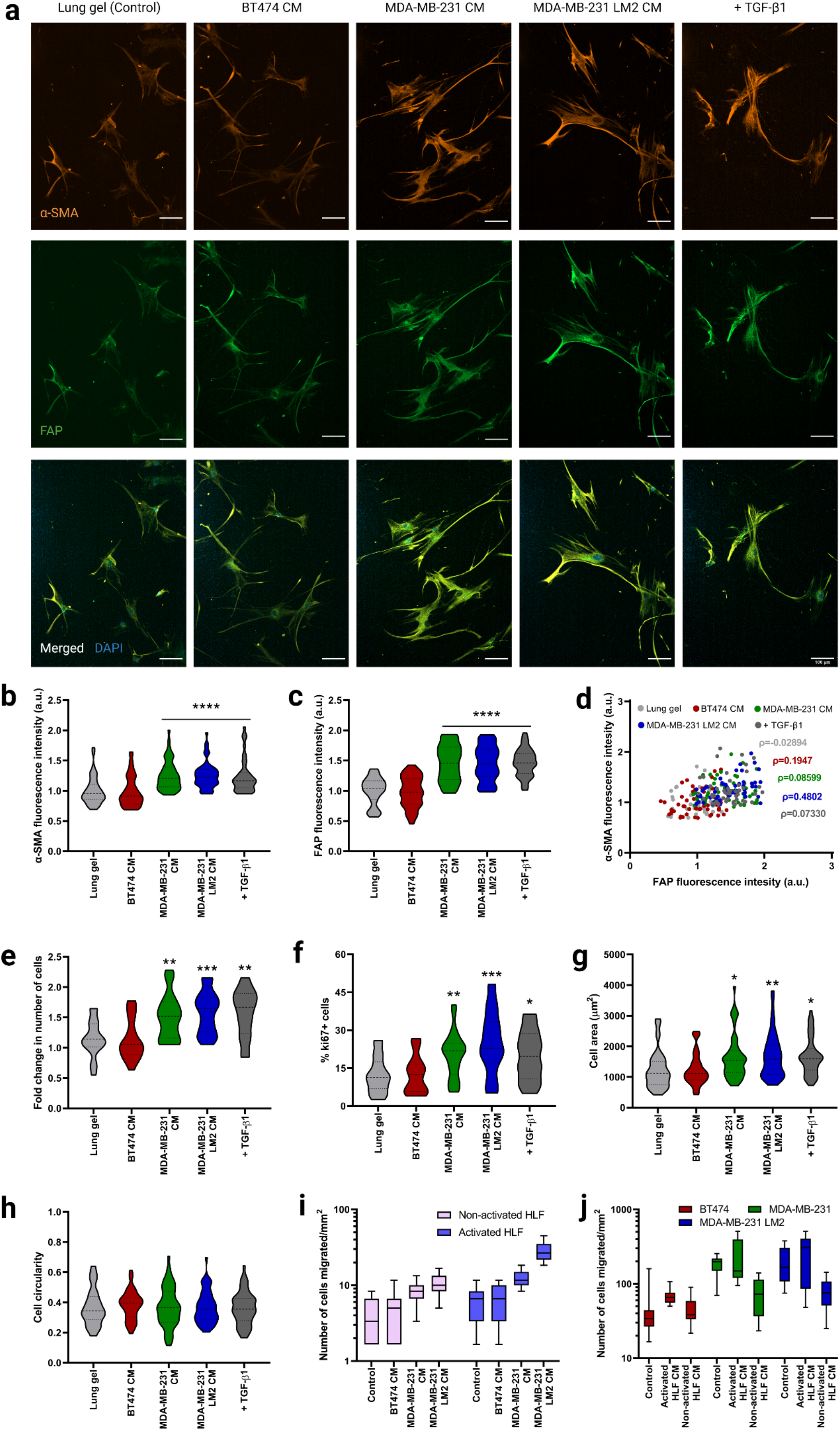
Fibroblast phenotype and activation in lung hydrogels in breast cancer conditioned media. a) Representative fluorescent images of HLFs cultured in 3D lung hydrogels with breast cancer conditioned media showing α-SMA (orange), and FAP (green) expressions along with merged HLF images with nuclei staining with DAPI (blue). Scale bar: 100 µm. b) Quantification of α-SMA expression from HLFs cultured in lung gels with breast cancer conditioned media MDA-MB-231 CM, MDA-MB-231 LM2 CM and BT474 CM along with controls. c) Quantification of FAP expression from HLFs cultured in lung gels with the different media conditions. d) Correlation between α-SMA and FAP expressions from HLFs cultured in lung gels with the different media. e) Fold change in cell count for HLFs cultured in lung gels with the different media conditions. f) Percentage of proliferative Ki67+ cells in HLFs cultured in lung gels with the different media conditions. g) Cell area for HLFs cultured in lung gels the different media conditions. h) Cell circularity for HLFs cultured in lung gels with the different media conditions. i) Quantification of number of non-activated and activated cells migrated towards breast cancer conditioned media in trans-well cell migration assay. j) Quantification of number of breast cancer cells migrated towards non-activated and activated HLF conditioned media. All data are mean + s.d. Statistical analyses were performed using Prism (GraphPad). Data in (b), (c), (e), (f), (g), (h), (i) and (j) were analyzed using a one-way analysis of variance followed by a Dunnett’s multiple comparison test with 95% confidence interval. *, **, ***, and **** indicate P < 0.05, P < 0.01, P < 0.001, and P < 0.0001.

TGF-β1 has been shown to activate fibroblasts and differentiate them into myofibroblasts^42,43^. TGF-β1 activates c-Jun-NH2-terminal kinase (JNK), mitogen-activated protein kinase (MAPK), and extracellular signal-regulated kinase (Erk) in fibroblasts^44–46^, and TGF-β1 activated JNK regulates the phenotypic modulation of HLFs to myofibroblasts^42^. We cultured HLFs in serum-free FBM for 6 days on TCPS and they were incubated with 20ng/mL TGF-β1 for 96 hours. TGF-β1 induced HLF activation within 96-hour incubation (Supplementary Figure S5a, c, d). Activated HLFs expressed significantly higher amounts of the fibroblast activation markers α-SMA (Supplementary Figure S5c), FAP (Supplementary Figure S5d), and Ki67 (Supplementary Figure S5e) compared to HLFs cultured without TGF-β1. Activated HLFs had a higher cell surface area than quiescent HLFs (Supplementary Figure S5f). We similarly cultured the HLFs in the 3D lung hydrogels, incubating them with TGF-β1 for 96 hours. Like the HLFs cultured on TCPS, HLFs exhibited an activated phenotype in the lung hydrogels (Figure 3). These results demonstrated reproducible TGF-β1 mediated activation of HLFs on both 2D control surfaces and in the 3D lung hydrogels.

### 2.4 HLF activation in 3D lung gels via metastatic breast cancer cells

The mechanism by which local, healthy lung fibroblasts remodel the microenvironment to create a favorable soil for metastasizing breast cancer cells is linked to HLF activation ^47^. As a potential application of our synthetic lung hydrogels, we quantified HLF activation in a metastatic niche with conditioned media collected from lung tropic MDA-MB-231 LM2 cells, the parental MDA-MB-231, and a non-metastatic control BT474 cell line. HLFs cultured in metastatic MDA-MB-231 LM2 CM and parental MDA-MB-231 CM for 6 days exhibited activated phenotypes and expressed significantly higher amounts of α-SMA and FAP (Figure 3a-d) and proliferated more (Figure 3e, f) than HLFs cultured in FBM. The phenotype of these cells was similar to the ones incubated with TGF-β1, validating the activation of the HLFs. On the other hand, HLFs cultured in non-metastatic BT474 CM maintained their quiescence, as observed from their activation marker expression (Figure 3a, b, c). The activated HLFs were significantly more spread than the non-activated cells as observed from their cell area (Figure 3g).

To test HLF’s migration potential in the metastatic niche we performed a transwell migration assay with HLFs and MDA-MB-231 LM2 CM, MDA-MB-231 CM, or BT474 CM. Both activated (+TGF-b) and quiescent HLFs (-TGF-b) migrated more towards the lung-tropic MDA-MB-231 CM and parental MDA-MB-231 CM compared to BT474 CM and FBM, with migration towards MDA-MB-231 LM2 CM significantly higher than towards MDA-MB-231 CM (Figure 3i). No significant difference was seen in HLF migration towards non-metastatic BT474 CM compared to the control. The activated HLFs exhibited significantly higher migration towards the metastatic conditions than the non-activated HLFs, while both activated and non-activated HLFs had similar invasion towards the non-metastatic and control conditions.

Conversely, we examined whether CM from HLFs stimulated increased invasion from metastatic breast cancer cells. We collected conditioned media from both activated HLF (activated HLF CM) and healthy non-activated HLF (non-activated HLF CM) cultures, with normal growth media as a control. Activated HLF CM induced higher breast cancer cell invasion than non-activated HLF CM for metastatic breast cancer cells, while non-activated HLF CM exhibited suppression of cancer cell invasion compared to the control medium (Figure 3j). Non-metastatic BT474 cells had similar responses to all the media treatment groups. Also, the metastatic MDA-MB-231 LM2 and MDA-MB-231 cells exhibited significantly higher migration than non-metastatic BT474 cells across treatments. No significant difference was seen in migration potential between the parental MDA-MB-231 cells and the lung tropic MDA-MB-231 LM2 cells.

### 2.5 TNC regulation of HLF activation in 3D lung gels

Finally, we aspired to demonstrate that the lung environment created here could be used to study cell-ECM interactions that are also observed *in vivo*. Again, we turned to metastatic breast cancer as a model for such a demonstration. First, we injected metastatic MDA-MB-231 LM2 CM, BT474 CM, or PBS into nude BALB/c female mice intraperitoneally for 10 consecutive days and harvested their lungs (Figure 4a). As expected, given work from Gocheva *et al*.^48^, the metastatic MDA-MB-231 LM2 CM induced upregulation of TNC expression in the lung compared to the non-metastatic BT474 CM and PBS (Figure 4b, c). A higher number of activated fibroblasts were observed in the lungs of the mice injected with MDA-MB-231 LM2 CM than those injected with BT474 CM and PBS, as represented by high fluorescence intensity of the activation marker α-SMA.

**Figure 4.**
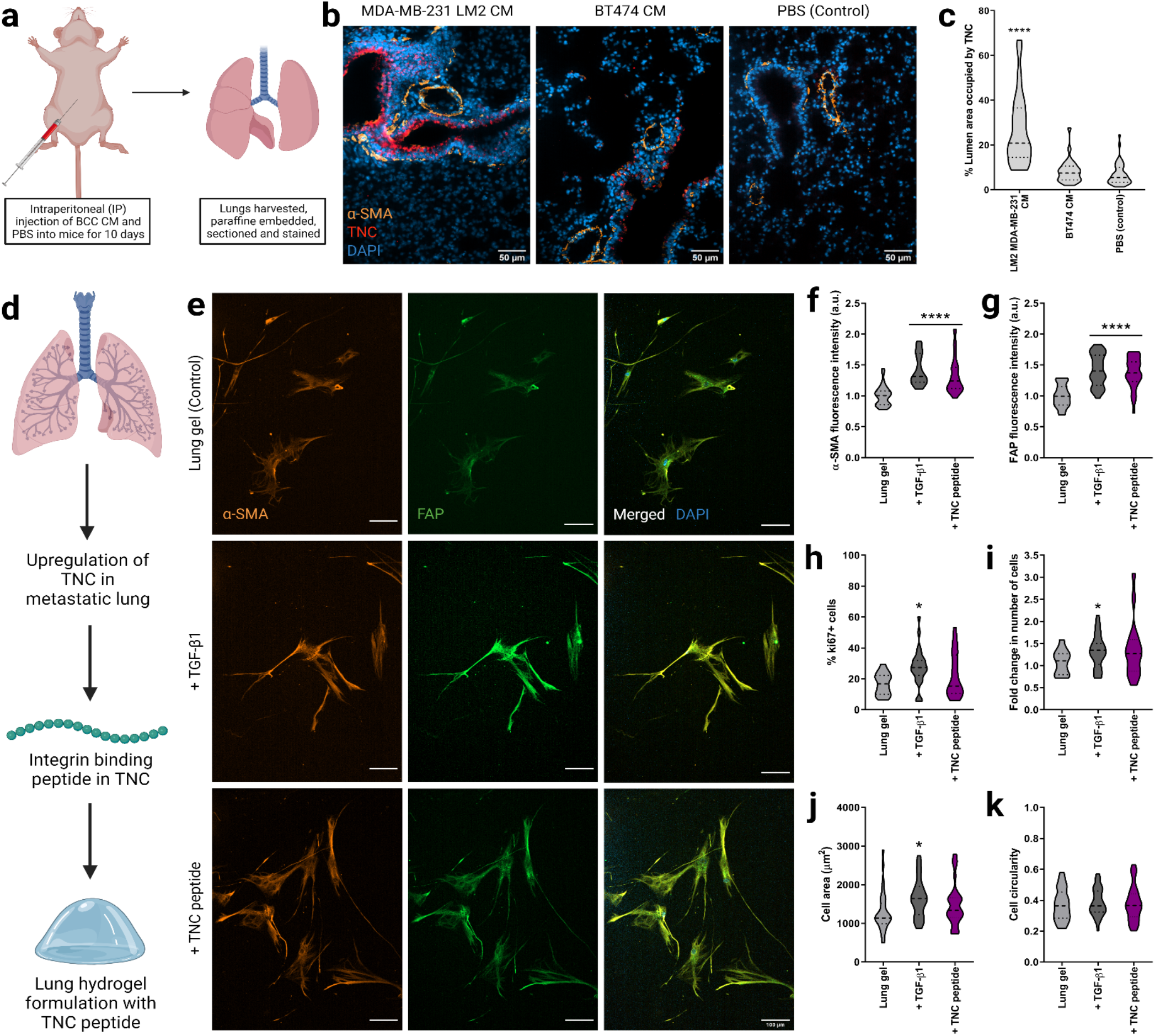
Fibroblast phenotype and activation in lung hydrogels with TNC peptide. a) Schematic of harvesting the lung tissues from nude BALB/c female mice injected with breast cancer conditioned media or PBS (control) for 10 days. b) Representative fluorescent images of lung sections with MDA-MB-231 LM2 CM, BT474 CM and PBS (control) showing DAPI staining for nuclei (blue), a-SMA (orange), and TNC (red) expressions along with merged images. Scale bar: 50 μm. c) Quantification of area coverage by TNC within the lung sections from mice injected with breast cancer conditioned media or PBS. d) Schematic of lung hydrogel design with TNC peptide. e) Representative fluorescent images of HLFs cultured in 3D lung hydrogels with TNC peptide showing a-SMA (orange), and FAP (green) expressions along with merged HLF images with nuclei staining with DAPI (blue). Scale bar: 50 μm. f) Quantification of a-SMA expression from HLFs cultured in lung gels with TNC peptide along with controls. g) Quantification of FAP expression from HLFs cultured in lung gels with TNC peptide along with controls. h) Percentage of proliferative ki67+ cells in HLFs cultured in lung gels with TNC peptide along with controls. i) Fold change in cell count for HLFs cultured in lung gels with TNC peptide, along with HLFs cultured in growth media with or without TGF-β1 as controls. j) Cell area for HLFs cultured in lung gels with TNC peptide along with controls. k) Cell circularity for HLFs cultured in lung gels with TNC peptide along with controls.tide cocktail. All data are mean + s.d. Statistical analyses were performed using Prism (GraphPad). Data in (a), (b), (c), (d), (g) and (h) were analyzed using a one-way analysis of variance followed by a Dunnett’s multiple comparison test with 95% confidence interval. *, **, ***, and **** indicate P < 0.05, P < 0.01, P < 0.001, and P < 0.0001.

We then used this *in vivo* finding to alter the lung hydrogel ECM design. We quantified the ECM of metastatic lung tissue by combining the healthy ECM composition presented earlier and the metastatic lung ECM quantification by Gocheva *et al*.^48^ We added representative peptide sequences for TNC to our peptide cocktails to mimic the metastatic ECM composition. The relative amounts of the integrin-binding and MMP-degradable peptides in the metastatic lung ECM mimetic MMP-degradable and integrin-binding peptide cocktails are presented in Supplementary Figure S6. We incorporated these metastatic peptide cocktails in our PEG-MAL gels to create a metastatic lung hydrogel platform.

We then encapsulated HLFs in the 3D metastatic lung hydrogels (TNC-containing) and cultured them in normal growth medium FBM for 6 days (Figure 4d). We cultured HLFs in healthy lung hydrogels with serum-free FBM and TGF-β1 for 96 hours as a positive control and with only FBM as a negative control. HLFs cultured in metastatic (TNC-containing) lung hydrogel exhibited an activated phenotype, with significantly higher amounts of α-SMA and FAP than HLFs cultured in healthy lung hydrogel (Figure 4e-g). The phenotype of these TNC-treated cells was similar to the cells incubated with TGF-β1, which confirmed the activation of the HLFs in the metastatic lung hydrogels. However, no significant difference in cell proliferation was observed for the HLFs cultured in TNC-containing metastatic lung hydrogels compared to the HLFs cultured in healthy lung hydrogels as observed from the fold-change in the number of cells and Ki67 expression (Figure 4h, i). Also, the morphology of TNC-activated cells was not significantly different compared to the quiescent cells in the healthy lung hydrogels as seen from their cell area and circularity (Figure 4j, k). Overall, this work demonstrates that our lung hydrogels support HLF quiescence, can be used to study HLF activation via known stimulation routes such as TGF-b, support HLF activation and motility from metastatic breast cancer cells, and have ECM plug-and-play capability to study how individual ECM components drive cell behaviors.

## Discussion

While cancer–ECM interactions are instrumental in the progression of cancer metastasis, the role of altered lung microenvironment on breast cancer cell survival and growth remains unclear. Scherzer *et al*. demonstrated that decellularized human fibroblast-derived ECM promoted the growth of lung adenocarcinoma cells^49^. Reports have shown that breast cancer cells that infiltrate the lungs can support the survival and growth of pulmonary micrometastases by expressing the ECM protein tenascin C (TNC)^50^. TNC has also been associated with poor metastasis-free survival in patients. Tumor cells prepare their own metastatic niche by producing ECM components and inducing activation of lung fibroblasts^51^. TNC has been identified as a promoter of both lung cancer progression and pulmonary metastasis formation^50,52,53^. However, very few studies have studied the effect of such a niche on the phenotype of the resident cells. Incorporation of the TNC integrin-binding peptide into our lung hydrogel induces HLF activation, further validating the involvement of TNC in facilitating breast-to-lung metastasis. We foresee the use of this synthetic lung hydrogel to understand extracellular mechanisms of fibroblast activation, as well as to identify individual contributions of quiescent and activated fibroblasts in healthy and diseased lung tissues respectively.

Studies have investigated breast cancer cell behavior in lung fibroblast-derived matrices, which recapitulates the *in vivo* proteomic landscape^54^. Multiple studies have explored the roles of cancer-associated fibroblasts (CAFs) and metastasis-associated fibroblasts (MAFs) on the migration of breast cancer cells. Compared with CAFs within primary tumors, MAFs have a stronger ability to enhance the proliferation, migration, and invasion potential of breast cancer cells^55,56^. MAFs facilitate the metastatic cancer cell invasive phenotype possibly via TGF-β1^57^.Studies have shown that fibroblasts are activated in stiffened, fibrotic, or metastatic microenvironments^15,47^. Fibroblast activation often leads to differentiation into myofibroblasts, characterized by prominent stress fiber formation containing α-SMA^14–19^ and high expression of FAP^58^. TGF-β1 aids in tissue remodeling and matrix stiffening by inducing increased production of ECM proteins, particularly fibronectin mediated by TGF-β1 activated JNK^59,60^. Conte *et al*. demonstrated that pirfenidone, an orally active small molecule that has been shown to inhibit the progression of fibrosis in idiopathic pulmonary fibrosis patients, reduces HLF proliferation and TGF-β mediated differentiation into myofibroblasts by attenuating key TGF-β induced signaling pathways^61^.

When activated via TGF-β1, fibroblasts show a characteristic biphasic response, with the peak migration velocities observed at nanomolar concentrations^62^. In the human body, activated fibroblasts show high migration potential at short time scales as prompt fibroblast migration is required for wound-healing purposes. Studies have reported increased fibroblast migration when cultured with TGF-β1 for a short time (6 hours), however, this migration potential was significantly reduced when fibroblasts were cultured with TGF-β1 for a longer period (6 days)^63^. Here we show that HLF activation in our lung hydrogel is induced by the pro-fibrotic cytokine TGF-β1 or by metastatic breast cancer conditioned media. We also show that injecting metastatic breast cancer conditioned media into nude BALB/c female mice promotes TNC deposition in the mouse lungs, which has been shown to aid in survival and outgrowth of pulmonary micrometastases.

Bioengineers have designed several *in vitro* lung ECM mimetic models^64–66^. However, there is a lack of tunable synthetic *in vitro* models that can mimic and independently control the mechanical and biochemical features of the lung ECM. Here, we combined proteomics, bioinformatics, and biomechanics to make a lung-mimetic PEG hydrogel. Integrins are the largest class of cell adhesion receptors that mediate attachment to the ECM and activate intracellular signaling^67^, and collectively the MMP family can degrade most proteins in the ECM^68^. Besides mimicking the lung modulus, this lung ECM hydrogel includes 17 different peptides to capture the diverse integrin-binding and MMP-sensitive domains of the lung ECM proteins. Having our synthetic, PEG-based hydrogel match the lung modulus is important because the modulus of the ECM regulates cell behavior and functions^69^. This work, along with our recent works to mimic brain and bone marrow^30,31^, demonstrates a novel and improved approach to biomimetic hydrogel design. The bi-directional interactions between lung ECM and resident fibroblasts, wherein the ECM provides biochemical and mechanical cues to maintain fibroblast phenotype and functions while fibroblasts deposit much of the ECM, play a major role in maintaining tissue homeostasis and tissue functions^1^. Using our 3D lung hydrogel, we can maintain quiescence of human lung fibroblasts (HLFs). Future studies investigating the effects of the cell-ECM interactions in driving tissue fate and disease progression using our lung ECM hydrogel will allow us to understand the complex signaling mechanisms underlying these interactions. This lung hydrogel can also be used as a high throughput platform for screening potential drug targets for various lung diseases.

## Materials and methods

### Cell culture

All cell culture supplies were purchased from Thermo Fisher Scientific (Waltham, MA, USA) unless otherwise noted. Human lung fibroblast (HLF) cells were purchased from American Type Culture Collection (ATCC) (Manassas, VA, USA) and cultured in Fibroblast basal medium (FBM) (Lonza, Quakertown, PA, USA) supplemented with 2% fetal bovine serum (FBS), 0.1% insulin, 0.1% r-human fibroblast growth factor (rhFGF) and 0.1% Gentamicin sulfate amphotericin-B (GA-1000) (Lonza, Quakertown, PA, USA). HLFs were used from passages 1 to 10. Human breast cancer cell line MDA-MB-231 was provided by Dr. Shannon Hughes. Lung tropic MDA-MB-231 LM2 and MDA-MB-231 cell lines were cultured in Dulbecco’s modified eagle’s medium (DMEM), supplemented with 1% penicillin-streptomycin, and 10% FBS. BT474 cell line was cultured in Roswell Park Memorial Institute (RPMI) 1640 medium, supplemented with 1% penicillin-streptomycin, and 10% FBS.

### Liquid chromatography-mass spectrometry (LC-MS/MS)

Six biological replicates were analyzed for human lung tissue samples. The ECM-rich pellet remaining from the CNCMS kit was solubilized and reduced in 8 M urea, 100 mM of ammonium bicarbonate, and 10 mM dithiothreitol (DTT) for 30 minutes at pH 8 and 37°C. Samples were alkylated with 25 mM iodoacetamide (Sigma-Aldrich) in the dark at room temperature for 30 minutes before the solution was quenched with 5 mM DTT. Before cleavage, the solution was diluted to 2 M urea with 100 mM ammonium bicarbonate at pH 8. Proteins were cleaved via trypsin and Lys-C endoproteinase (Promega, Madison, WI), at a ratio of 1:50 enzyme to protein overnight (12-16 hours) at 37°C. Samples were cleaned and concentrated using a C18 column. A reverse phase LC gradient was used to separate peptides before mass analysis. Mass spectrometry analysis was performed in an Orbitrap Fusion Tribrid. Peptides were aligned against the Matrisome using the Thermo Proteome Discoverer 1.41.14^70^. Parameters used: trypsin as a protease, with 4 missed cleavage per peptide, a precursor mass tolerance of 10 ppm, and fragment tolerance of 0.6 Da.

### Identifying integrin-binding and MMP-degradable proteins in lung

The lung ECM composition was assessed via histology-based bioinformatics and proteomics datasets. Histology data from the Human Protein Atlas was used to identify lung proteins, their expression, and local distribution. A scoring system was based on the number of unique peptides, and peptide spectrum matches (PSM) obtained from the LC-MS/MS data and relative protein detection level (not detected or ND, low, medium, or high) to quantify and screen the lung ECM proteins. From this list of ECM proteins, those with integrin-binding capabilities and susceptibility to matrix metalloproteinase (MMP) were chosen to be incorporated into the hydrogel to allow for cell adhesion and synthetic matrix degradation. Bioactive motifs of the selected proteins previously identified in the literature were synthesized and incorporated into the hydrogel to mimic the proteomic landscape of the lung ECM. The scoring system was also used to determine the relative amounts of each integrin-binding peptide and MMP-degradable crosslinker.

### Solid-phase peptide synthesis

All peptides were synthesized on a CEM Liberty Blue automated solid phase peptide synthesizer (CEM, Mathews, NC) using Fmoc protected amino acids (Iris Biotech GMBH, Germany). The peptide was cleaved from the resin by sparging-nitrogen gas through a solution of peptide-resin and trifluoroacetic acid (TFA), triisopropylsilane, water, and 2,2′-(Ethylenedioxy)diethanethiol at a ratio of 92.5:2.5:2.5:2.5 % by volume, respectively (Sigma-Aldrich, St. Louis, MO) for 3 hours at room temperature in a peptide synthesis vessel (ChemGlass, Vineland, NJ). The peptide solution was filtered to remove the resin and the peptide was precipitated out using diethyl ether at -80°C. Molecular mass was validated using a MicroFlex MALDI-TOF (Bruker, Billerica, MA) using α-cyano-4-hydroxycinnamic acid as the matrix (Sigma-Aldrich). Peptides were purified to ≥ 95% on a VYDAC reversed-phase c18 column attached to a Waters 2487 dual **λ** adsorbable detector and 1525 binary HPLC pump (Waters, Milford, MA).

The following sequences were synthesized: GCGFYFDLR, GCGRKRK, GPRGGC, GCGWTVFQKRLDGS, CGPHSRNGGGGGGRGDS, GCKQLREQ, GCRDSGESPAYYTADRCG, GCRDVPLSLTMGDRCG,

CSRARKQAASIKVAVADR, GCRDVPMSMRGGDRCG, GCRDRPFSMIMGDRCG, GCRDVPLSLYSGDRCG,

GCRDGPLGLWARDRCG, and GCRDIPESLRAGDRCG.

The following sequences were purchased from GenScript (Piscataway, NJ, USA) at ≥ 95% purity: CGP(GPP)5GFOGER(GPP)5, CGP(GPP)5GFOGER(GPP)5, GRGDSPCG.

### Synthesis of 3D lung hydrogels

10 mM of 10 kDa 4-arm PEG-maleimide (Jenkem Technology, Plano, TX) was reacted with 2mM of the lung integrin-binding peptide cocktail for 10 minutes in 0.5X PBS at pH 7.4. This solution was crosslinked at a 1:1 molar ratio of thiol to maleimide in 0.5X PBS at pH 7.4, yielding 10 wt% of polymer and peptide in the synthesis mixture, and the crosslinker cocktail was composed of 75 molar% of 1.5 kDa linear PEG-dithiol (Jenkem) and 25 molar% of the MMP-degradable cocktail. Gels were polymerized in 10 μL volumes with 5,000 cells/μL, and cell culture medium was added after 5 minutes to swell the material for at least 18 hours before use.

### Hydrogel mechanical and structural characterization

The effective Young’s modulus was measured using indentation testing on 10 μL volumes of the 3D hydrogels. A custom-built instrument was used as previously described^71^. Lung mechanical data was taken from Polio *et al*.^39^ For this application, a flat punch probe with a radius of 0.5 mm was applied to samples at a fixed displacement rate of 10 μm/s, for a maximum displacement of 300 μm. The first 10% of the linear region of the force-indentation curves was analyzed using a Hertzian model modified by Hutchens *et al*. to account for dimensional confinement described by the ratio between the contact radius (*a*) and the sample height (*h*) (0.5<*a*/*h*<2)^72^. The force (*F*) and displacement (*δ*) measurements were recorded with a custom LabVIEW program (National Instruments, Austin, TX, USA). The material compliance (*C*) was measured over the linear region after contacting the surface using the following equation:

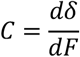

The deflection of the cantilever was addressed by subtracting the deflection required to produce the given force from the displacement indicated. Measurements were conducted at 2 or 3 different locations per sample away from the edge of the sample. The Young’s modulus (*E*) was calculated using the following equation after taking the height of the sample (*h*), the Poisson’s ratio of 0.42 (*ν*), and the radius of the circular indenter (*a*) into account, assuming the material is elastic, according to Shull *et al*.136.

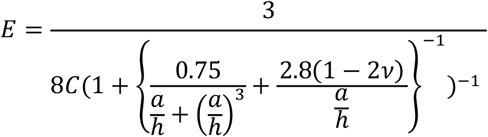

### Cell attachment assay

Glass coverslips were prepared with 1 ug/cm^2^ of the lung peptide coupled to the surface using silane chemistry described by Barney *et al*^40^. Cells were seeded at 3,000 cells/cm^2^ on functionalized glass coverslip surfaces containing individual integrin-binding peptides at the same molar concentrations as in the lung integrin-binding peptide cocktail or the entire lung integrin-binding peptide cocktail at 1 µg/cm^2^ concentration. The HLFs were seeded in serum-free FBM (Lonza, Quakertown, PA, USA), and MDA-MB-231 LM2 cells were seeded in serum-free DMEM. The cells were imaged in an environment-controlled Zeiss Axio Observer Z1 microscope (Carl Zeiss, Oberkochen, Germany) using an AxioCam MRm camera and an EC Plan-Neofluar 20X 0.4 NA air objective. Images were taken every five minutes for an incubation period of 2 hours. Cells were traced and cell area was assessed over time in ImageJ (NIH, Bethesda, MD, USA).

### Competitive binding assay

Glass coverslips were prepared with 1 ug/cm^2^ of the lung peptide coupled to the surface using silane chemistry described by Barney *et al*^40^. Cells were seeded at 3,000 cells/cm2 in their normal growth medium without serum after 30 minutes of pre-treatment with individual peptides or the complete lung integrin-binding peptide cocktail. Lung integrin-binding peptide cocktail was dosed at a molar amount of 400 nmol/mL of medium and the molar amount dosed for each individual peptide was as follows: GRGDSPCG at 164 nmol/mL, GCGWTVFQKRLDGS at 52nmol/mL, GPRGGC at 44 nmol/mL, CGP(GPP)5GFOGER(GPP)5 and GCKQLREQ at 32 nmol/mL, CGPHSRNGGGGGGRGDS at 24 nmol/mL, GCGFYFDLR at 20 nmol/mL, and GCGRKRK, CGP(GPP)5GROGER(GPP)5, and CSRARKQAASIKVAVADR at 12 nmol/mL. The cells were imaged in an environment-controlled Zeiss Axio Observer Z1 microscope (Carl Zeiss, Oberkochen, Germany) using an AxioCam MRm camera and an EC Plan-Neofluar 20X 0.4 NA air objective. Images were taken every five minutes for an incubation period of 2 hours. Cells were traced and cell area was assessed over time in ImageJ (NIH, Bethesda, MD, USA).

### Cell attachment assay with cilengitide

Glass coverslips were prepared with 1 ug/cm^2^ of the lung peptide coupled to the surface using silane chemistry described by Barney *et al*^40^. Cells were seeded at 3,000 cells/cm^2^ in their normal growth medium without serum after 30 minutes of pre-treatment with cilengitide (Apex Biotechnology, Houston, TX, USA). Cilengitide was dosed at a molar concentration of 100 nmol/mL. The cells were imaged in an environment-controlled Zeiss Axio Observer Z1 microscope (Carl Zeiss, Oberkochen, Germany) using an AxioCam MRm camera and an EC Plan-Neofluar 20X 0.4 NA air objective. Images were taken every five minutes for an incubation period of 2 hours. Cells were traced and cell area was assessed over time in ImageJ (NIH, Bethesda, MD, USA).

### Quantitative reverse transcription polymerase chain reaction (qRT-PCR)

Cells were seeded at 250,000 cells per well in a 6-well plate and allowed to grow to form a uniform monolayer. Cells were then washed with PBS and lysed. Total RNA was extracted using the Genelute Mammalian Total RNA kit (Sigma-Aldrich) followed by cDNA synthesis using the RevertAid reverse transcriptase protocol, with the exception of using RNasin 40 U/mL (Promega, Madison, WI) as the RNase inhibitor. cDNA was then amplified using specific primers and the Maxima Sybr green master mix (Thermo Fisher Scientific) on a Rotor-Gene Q thermocycler (Qiagen, Valencia, CA) for 45 cycles Both β-actin and ribosomal protein S13 were included as reference genes to permit gene expression analysis using the 2-ddCt method.

### Conditioned media collection

Breast cancer conditioned media were collected from metastatic breast cancer cells, parental MDA-MB-231 and lung tropic MDA-MB-231 LM2, and non-metastatic breast cancer cells BT474. Fibroblast-conditioned media were collected from healthy HLFs and activated HLFs. Breast cancer cells were cultured in their respective growth media in T75 cell culture flasks until cells were 70-75% confluent. The cells were then cultured in serum-free media and the media were subsequently collected after 48 hours. HLFs were similarly cultured in FBM until 75-80% confluence. HLFs were then cultured in serum-free FBM for 48 hours and the medium was collected. For culturing activated HLFs, cells were cultured in serum-free FBM, supplemented with TGF-β1 (Fisher Scientific, Hampton, NH, USA) at a final concentration of 20 ng/mL. The activated conditioned medium was collected when the cells are 80-85% confluent. The conditioned medium was filtered through a 0.45 µm syringe filter and stored at -80°C until use.

### HLF activation assay

HLF activation assays were performed both on 2D and 3D. For the 2D HLF activation assay, 5,000 cells were seeded per well in glass bottom 24 well plate serum-free FBM. HLFs were incubated with solubilized TGF-β1 at a final concentration of 20 ng/mL for 96 hours to induce activation of the HLFs. The cells were then fixed and stained for α-SMA, FAP, Ki67, and DAPI. The stained cells were subsequently imaged on the Zeiss Spinning Disc Observer Z1 microscope (Carl Zeiss, Oberkochen, Germany).

For the 3D HLF activation assay, 5,000 cells were encapsulated in each lung hydrogel, which was an optimized HLF cell density in the lung hydrogel to prevent cell clustering over long culture periods. HLFs were cultured in SFM incubated with solubilized TGF-β1 at a final concentration of 20 ng/mL or in breast cancer conditioned media supplemented with FBS for 96 hours to induce activation of the HLFs. The cells were then fixed and stained for α-SMA, FAP, and Ki67. The stained cells were subsequently imaged on the Zeiss Spinning Disc Observer Z1 microscope (Carl Zeiss).

### Cell migration/invasion assay

HLFs and activated HLFs were seeded into the top of a Boyden chamber in serum-free FBM, at a density of 50,000 cells/well. The bottom chamber was filled with breast cancer conditioned media or normal growth media, both supplemented with FBS. Cells were allowed to invade through 8 µm pores for 48 hours. Then the cells on top of the Boyden chamber were removed. The bottom of the Boyden chamber and the bottom chamber i.e., the well were washed with 1X PBS and fixed in 4% formaldehyde. Cell nuclei were stained with DAPI, and the well and the bottom side of the Boyden chamber were imaged on the Zeiss Spinning Disc Observer Z1 microscope (Carl Zeiss) for manual quantification of invading cells.

Breast cancer cells (MDA-MB-231, MDA-MB-231 LM2, and BT474) were seeded into the top of a Boyden chamber in serum-free media, at a density of 100,000 cells/well. The bottom chamber was filled with HLF CM, activated HLF CM, or normal growth media, each supplemented with FBS. Cells were allowed to invade through 8 µm pores for 48 hours. Then the cells on top of the Boyden chamber were removed. The bottom of the Boyden chamber and the bottom chamber (i.e., the well) were washed with 1X PBS and fixed in 4% formaldehyde. Cell nuclei were stained with DAPI, and the well and the bottom side of the Boyden chamber were imaged on the Zeiss Spinning Disc Observer Z1 microscope (Carl Zeiss) for manual quantification of invading cells.

### Immunofluorescent staining

Cells were fixed at 6 days after seeding in 4% formaldehyde. Cells were then permeabilized in Tris-Buffered Saline (TBS) with 0.25% Triton-X for 10 minutes and washed 3 times in TBS with 0.1% Triton-X (TBS-T). Blocking was done for 30 minutes at room temperature in AbDil or TBS-T + 2 w/v% Bovine serum albumin (BSA, Sigma Aldrich) for 2D and in TBS-T + 5 w/v% BSA for 3D. Cells were then incubated overnight in primary antibody in the blocking solution at 4°C using 1 or more of the following antibodies: α-SMA antibody (ab7817, 1:500, Abcam, Cambridge, MA, USA), FAP antibody (ab53066, 1:500, Abcam), Ki67 antibody (ab156956, 1:1000, Abcam. Cells were then washed 3 times in TBS-T and subsequently incubated in secondary antibody in AbDil for 1 hour for 2D and 2 hours for 3D at room temperature using the following antibodies: goat anti-mouse 555 (A21422, 1:200, Thermo Fisher Scientific), goat anti-rabbit 488 (A11070, 1:500, Thermo Fisher Scientific), or goat anti-rat 647 (A21247, 1:200, Thermo Fisher Scientific). Cells were again washed 3 times in TBS-T. Cells were then treated with DAPI (Vector Labs) at a 1:5,000 dilution for 5 minutes and washed in PBS 3 times. Cells were then imaged on the Zeiss Spinning Disc Observer Z1 microscope (Carl Zeiss).

### Intraperitoneal (IP) injections in BALB/c nude mice

The mice used for this study were 6 to 20 weeks old immunodeficient BALB/c nude (CAnN.Cg-Foxn1nu/Crl) female mice, ordered from Charles River Laboratories (Wilmington, MA, USA). Each mouse was weighed before injection. Breast cancer conditioned media MDA-MB-231 LM2 CM, BT474 CM, or 1X PBS were injected into the mice for 10 days. 5-6 mice were used for each of the media conditions. 150 µL of the pre-warmed media were injected into the animal’s lower right quadrant of the abdomen, to avoid damage to the urinary bladder, cecum, and other abdominal organs. The mice were sacrificed on day 11 and the brain, lungs, liver, and bones were harvested and fixed in 4% formaldehyde and embedded in paraffin before slicing them into 7 µm sections for immunohistochemistry. The protocol was approved by the Institutional Animal Care and Use Committee (IACUC).

### Immunohistochemistry

Immunohistochemistry was performed on the formalin fixed paraffin embedded (FFPE) mouse lung sections. Tissue sections were deparaffinized in xylenes with 2 washes for 10 minutes each (Fisher Scientific), rehydrated in ethanol with sequential two 10-minute washes with 100% ethanol (Fisher Scientific), one 5-minute wash with 95% ethanol (Fisher Scientific), one 5-minute wash with 70% ethanol and one 5-minute wash with 50% ethanol. The sections were then rinsed with deionized water and washed with 1X PBS for 10 minutes. Antigen retrieval was performed at 90°C for 15 minutes in Tris/EDTA buffer (pH 9.0), and then samples were cooled to room temperature for 30 minutes. The samples were then washed twice with TBS + 0.025% Triton X-100 for 5 minutes each. Non-specific protein absorption was blocked using two 1-hour incubations with Intercept Blocking Solution. Then the samples were incubated with primary TNC antibody (ab66045, 1:100, Abcam) and α-SMA antibody (ab7817, 1:500, Abcam) overnight at 4°C. Samples were then rinsed 3 times for 10 minutes each in 1X PBS, and incubated with the following antibodies for 1 hour at room temperature: goat anti-mouse 555 (A21422, 1:200, Thermo Fisher Scientific), or goat anti-rabbit 647 (A21244, 1:200, Thermo Fisher Scientific). The samples were then rinsed 3 times for 10 minutes each with PBS and were subsequently treated with DAPI (Vector Labs) at a 1:5,000 dilution for 5 minutes and washed in PBS 3 times. The samples were then mounted with GelVatol mounting medium (prepared in-house), and coverslips sealed with clear nail polish. Tissue sections were then imaged on the Zeiss Spinning Disc Observer Z1 microscope (Carl Zeiss).

### Statistical Analysis

Statistical analysis was performed with GraphPad Prism (9.0a) (GraphPad Software, Inc., La Jolla, CA, USA). Data are reported as the mean ± standard error. Unless otherwise noted, statistical significance was evaluated using a one-way analysis of variance (ANOVA). For the analysis, p-values <0.05 are considered significant, where p < 0.05 is denoted with *, < 0.01 with **, < 0.001 with ***, and < 0.0001 with ****.

## Acknowledgments

We would like to thank Dr. Sarah Perry, Dr. Alfred Crosby, and Dr. Lila Gierasch for their technical assistance and use of equipment. We thank Dr. Stephen Eyles and the Mass Spectrometry Core Facility at the Institute of Applied Life Sciences for support with LC-MS/MS. Research reported in this publication was supported by the Office of the Director, National Institutes of Health under Award Number S10OD010645. ANK was supported by an NIH Biotech Training Program (BTP) grant (T32 GM135096). CED was supported by an NIH Chemistry-Biology Interface (CBI) grant (T32 GM008515). SRP is a Pew Biomedical Scholar supported by the Pew Charitable Trusts. SRP was supported by a faculty development award from Barry and Afsaneh Siadat. This work was funded by an NIH New Innovator award (1DP2CA186573-01) and a National Science Foundation (NSF) CAREER grant (DMR-1454806) to SRP.

## Data Availability Statement

The raw/processed data required to reproduce these findings cannot be shared at this time due to technical or time limitations.

---

**Supplementary Figure 1.**
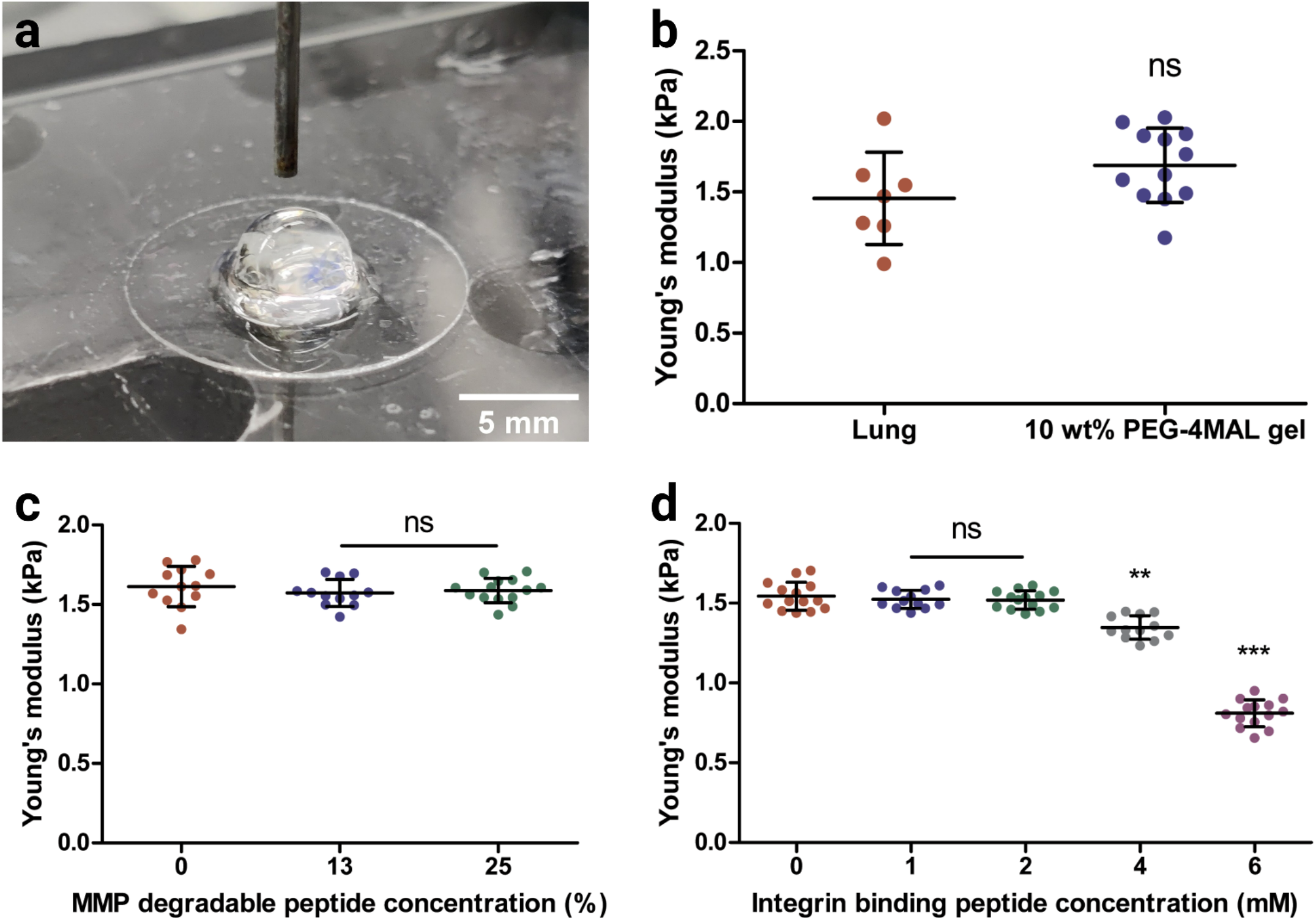
Mechanical characterization of lung hydrogels using micro-indentation method. a) Representative image of lung hydrogel after 24 hours of swelling in 1X PBS. Modulus of the hydrogel was measured using micro-indentation method with a 1 mm diameter flat probe. b) Effective modulus of 1O wt% 4-arm PEG-maleimide hydrogel without bioactive peptide ligands using micro-indentation replicated the lung modulus measured using the same method. c) Effective modulus of 10 wt% 4-arm PEG-maleimide hydrogel containing different molar concentrations of MMP degradable peptide crosslinkers using micro-indentation method. The tested peptide crosslinker molar concentrations were 0%, 13% and 25%. d) Effective modulus of 10 wt% 4-arm PEG-maleimide hydrogel containing different molar concentrations of integrin-binding peptide moieties using micro-indentation method. The tested peptide molar concentrations were O mM, 1 mM, 2 mM, 4 mM, and 6 mM.

**Supplementary Figure 2.**
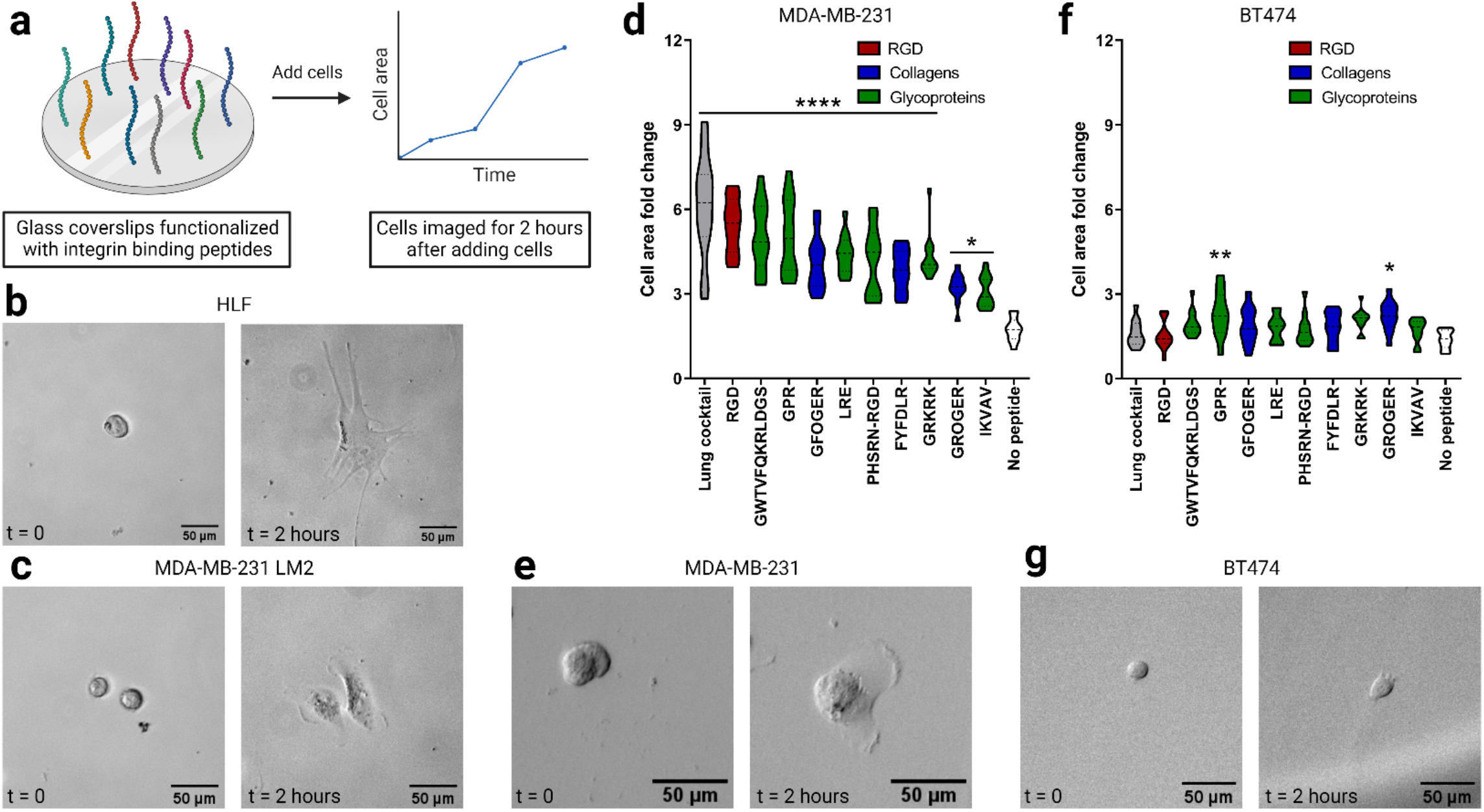
Functional validation of lung integrin-binding peptides. a) Schematics of the cell attachment assay where cells were seeded on glass coverslips functionalized with lung integrin-binding peptides and the cells were imaged for 2 hours. b) Representative images of HLFs at time t = 0 and t = 2 hours. c) Representative images of MDA MB-231 LM2 cells at time t = 0 and t = 2 hours. d) Cell area fold change for MDA-MB-231 cells in compared to negative control. e) Representative images of MDA-MB-231 cells at time t = 0 and t = 2 hours. f) Cell area fold change for BT474 cells in compared to negative control. g) Representative images of BT474 cells at time t = O and t = 2 hours.

**Supplementary Figure 3.**
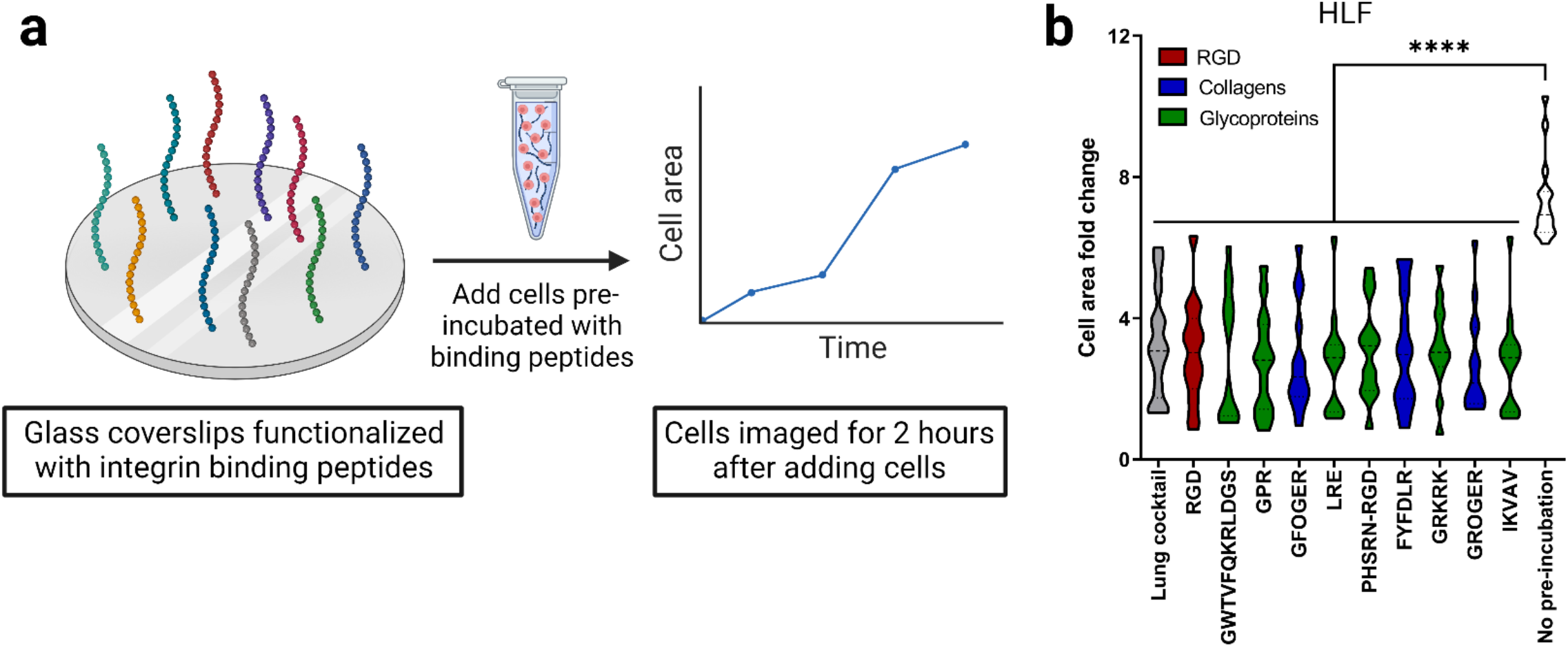
Competitive cell attachment assay on lung integrin-binding peptides using HLFs. a) Schematics of the competitive cell binding assay where HLFs pre-incubated with individual lung integrin-binding peptides were seeded on glass coverslips functionalized with lung integrin-binding peptide cocktail and the cells were imaged for 2 hours. b) Representative images of HLFs (pre-incubated with individual binding peptides) at time t = 0 and t = 2 hours.

**Supplementary Figure 4.**
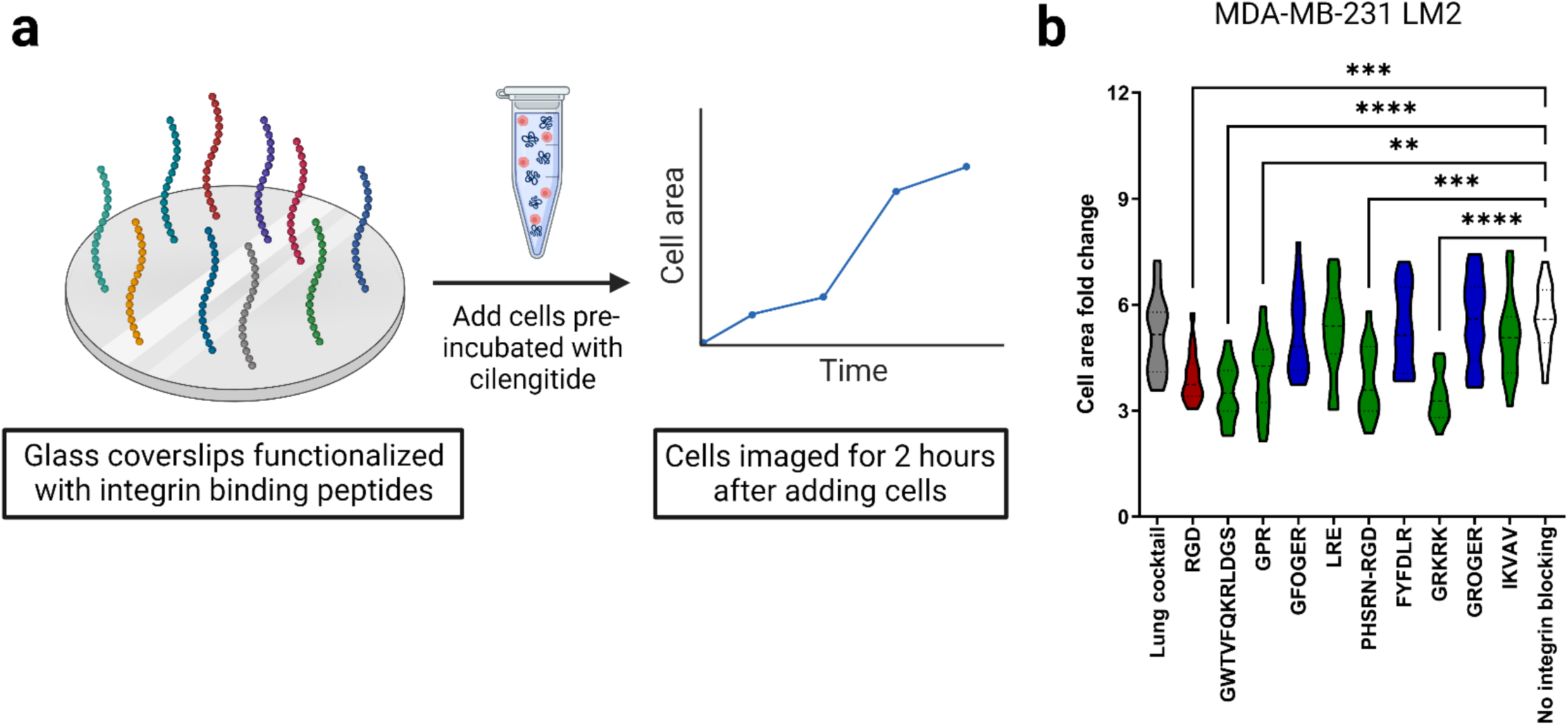
lntegrin-specific cell attachment with lung integrin-binding peptides. a) Schematics of the competitive cell binding assay where HLFs pre-incubated with cilengitide were seeded on glass coverslips functionalized with lung integrin-binding peptide cocktail and the cells were imaged for 2 hours. b) Cell area fold change 2 hours after seeding HLFs (pre-incubated with cilengitide) onto glass coverslips functionalized with integrin-binding peptide cocktail relative to a negative control (cells not pre-incubated with cilengitide).

**Supplementary Figure 5.**
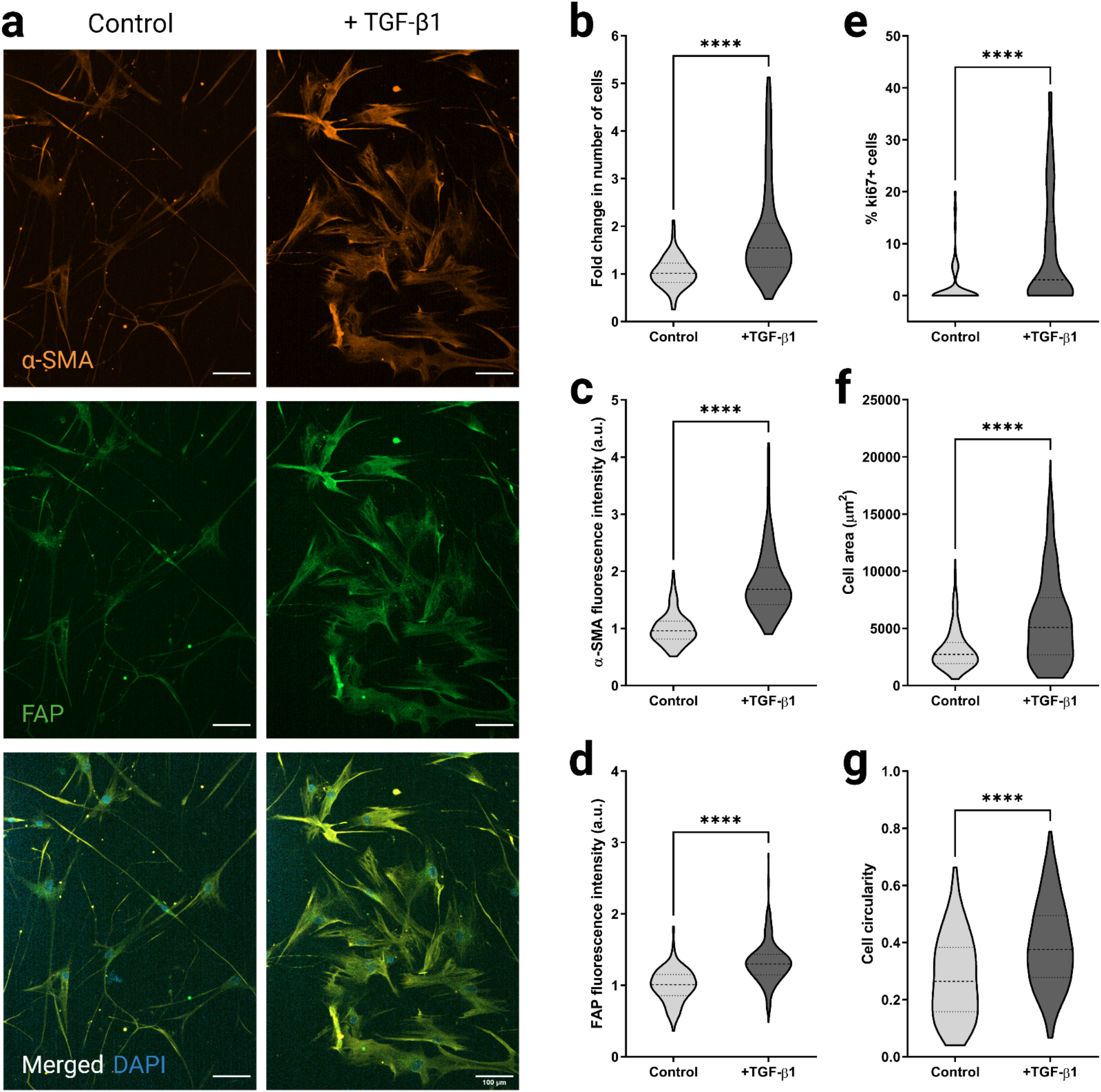
Fibroblast phenotype and activation on TCPS. (a) Representative fluorescent images of HLFs cultured on TCPS with or without pro-flbrotic cytokine, TGF-131 showing a-SMA (orange), and FAP (green) expressions along with merged HLF images with nuclei staining with DAPI (blue). (b) Fold change in cell count for non-activated (cultured without TGF-131) and activated (cultured with TGF-131) HLFs on TCPS representing cell proliferation characteristics. (c) Quantification of a-SMA expression from non-activated and activated HLFs cultured on TCPS. (d) Quantification of FAP expression from non-activated and activated HLFs cultured on TCPS. (e) Percentage of proliferative ki67+ cells in non-activated and activated HLF cultures on TCPS. (f) Cell area for non-activated and activated HLFs cultured on TCPS. (g) Cell circularity for non-activated and activated HLFs cultured on TCPS.

**Supplementary Figure 6.**
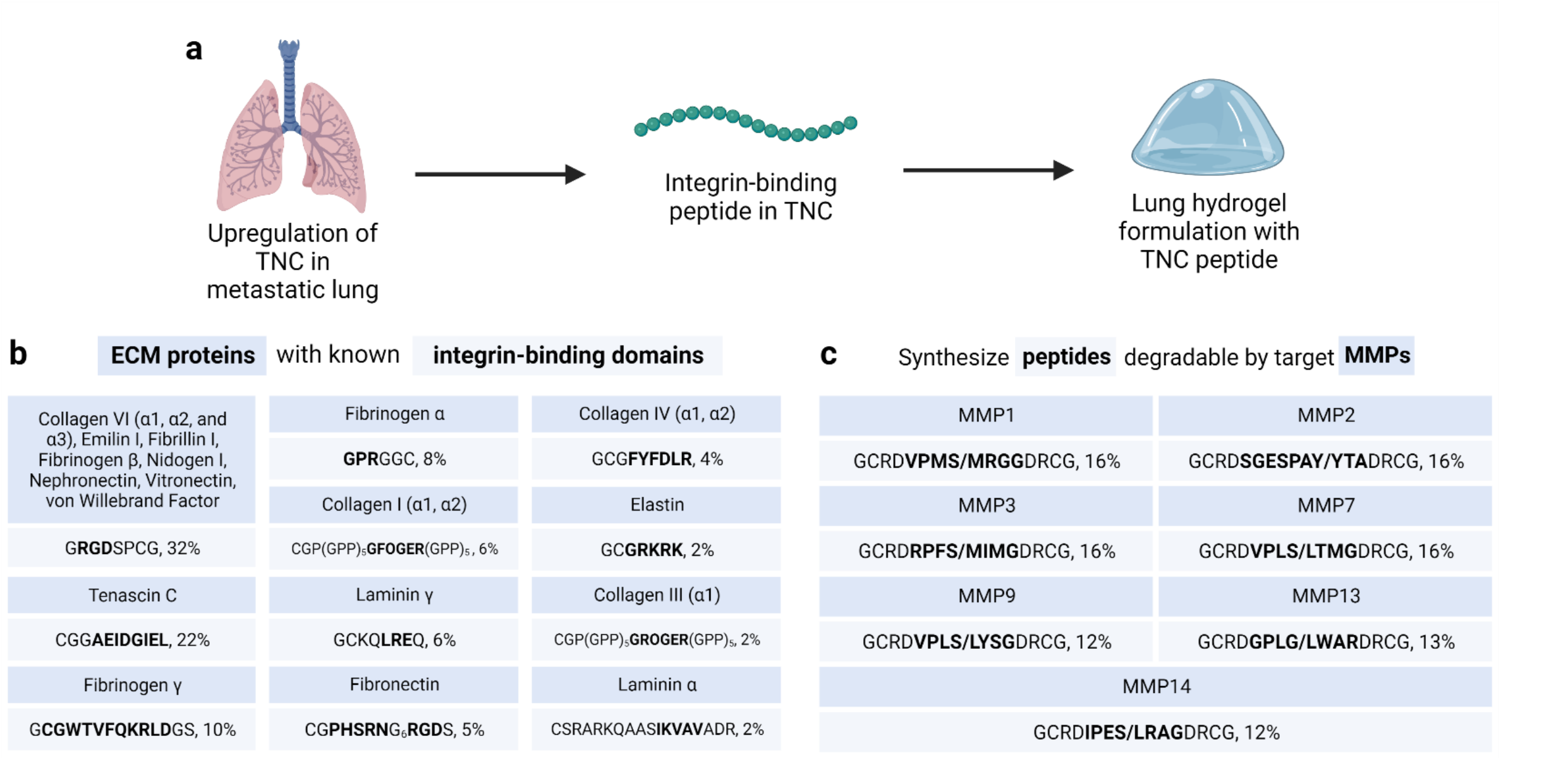
List and quantities of metastatic lung-specific integrin-binding and MMP-degradable peptides. a) Schematic of identification of metastatic lung ECM-specific integrin-binding and MMP-degradable peptides. b) List of identified integrin-binding peptide domains with their relative quantities corresponding to metastatic lung ECM proteins. c) List of identified MMP-degradable peptide domains with their relative quantities corresponding to the different MMPs.

## References

1. Dunsmore, S. E. & Rannels, D. E. Extracellular matrix biology in the lung. Am J Physiol 270, L3–27 (1996).

2. Grinnell, F. Fibroblast biology in three-dimensional collagen matrices. Trends Cell Biol 13, 264–269 (2003).

3. Du, P. et al. Human lung fibroblast-derived matrix facilitates vascular morphogenesis in 3D environment and enhances skin wound healing. Acta Biomater 54, 333–344 (2017).

4. Darby, I. A., Laverdet, B., Bonté, F. & Desmoulière, Fibroblasts and myofibroblasts in wound healing. Clinical, Cosmetic and Investigational Dermatology vol. 7 301–311 Preprint at https://doi.org/10.2147/CCID.S50046 (2014).

5. Kanta, J. Collagen matrix as a tool in studying fibroblastic cell behavior. Cell Adh Migr 9, 308–316 (2015).

6. Liu, S. et al. Expression of integrin β1 by fibroblasts is required for tissue repair in vivo. Journal of Cell Science vol. 123 3674–3682 Preprint at https://doi.org/10.1242/jcs.070672 (2010).

7. Zhao, X. K. et al. Focal adhesion kinase regulates fibroblast migration via integrin beta-1 and plays a central role in fibrosis. Sci Rep 6, 1–12 (2016).

8. Leask, A. Integrin β 1: A Mechanosignaling Sensor Essential for Connective Tissue Deposition by Fibroblasts. Adv Wound Care (New Rochelle) 2, 160–166 (2013).

9. White, E. S. Lung extracellular matrix and fibroblast function. Ann Am Thorac Soc 12, S30–S33 (2015).

10. Huang, X. et al. Matrix stiffness-induced myofibroblast differentiation is mediated by intrinsic mechanotransduction. Am J Respir Cell Mol Biol 47, 340–348 (2012).

11. Rhee, S. Fibroblasts in three dimensional matrices: Cell migration and matrix remodeling. Exp Mol Med 41, 858–865 (2009).

12. Eckes, B. et al. Mechanical tension and integrin α2β1 regulate fibroblast functions. Journal of Investigative Dermatology Symposium Proceedings vol. 11 66–72 Preprint at https://doi.org/10.1038/sj.jidsymp.5650003 (2006).

13. Kessler, D. et al. Fibroblasts in Mechanically Stressed Collagen Lattices Assume a ‘Synthetic’ Phenotype. Journal of Biological Chemistry 276, 36575–36585 (2001).

14. Asano, S. et al. Matrix stiffness regulates migration of human lung fibroblasts. Physiol Rep 5, 1–11 (2017).

15. Li, Z. et al. Transforming growth factor-β and substrate stiffness regulate portal fibroblast activation in culture. Hepatology vol. 46 1246–1256 Preprint at https://doi.org/10.1002/hep.21792 (2007).

16. Shinde, A. v., Humeres, C. & Frangogiannis, N. G. The role of α-smooth muscle actin in fibroblast-mediated matrix contraction and remodeling. Biochimica et Biophysica Acta - Molecular Basis of Disease vol. 1863 298–309 Preprint at https://doi.org/10.1016/j.bbadis.2016.11.006 (2017).

17. Hinz, B., Celetta, G., Tomasek, J. J., Gabbiani, G. & Chaponnier, C. Alpha-smooth muscle actin expression upregulates fibroblast contractile activity. Mol Biol Cell 12, 2730–2741 (2001).

18. Bharath Rao, K., Malathi, N., Narashiman, S. & Rajan, S. T. Evaluation of myofibroblasts by expression of alpha smooth muscle actin: A marker in fibrosis, dysplasia and carcinoma. Journal of Clinical and Diagnostic Research 8, 14–17 (2014).

19. Sapudom, J. et al. The interplay of fibronectin functionalization and TGF-β1 presence on fibroblast proliferation, differentiation and migration in 3D matrices. Biomaterials Science vol. 3 1291–1301 Preprint at https://doi.org/10.1039/c5bm00140d (2015).

20. Parker, M. W. et al. Fibrotic extracellular matrix activates a profibrotic positive feedback loop. Journal of Clinical Investigation vol. 124 1622–1635 Preprint at https://doi.org/10.1172/JCI71386 (2014).

21. Liu, F. et al. Feedback amplification of fibrosis through matrix stiffening and COX-2 suppression. Journal of Cell Biology 190, 693–706 (2010).

22. Kulasekaran, P. et al. Endothelin-1 and transforming growth factor-β1 independently induce fibroblast resistance to apoptosis via AKT activation. Am J Respir Cell Mol Biol 41, 484–493 (2009).

23. Horowitz, J. C. et al. Combinatorial activation of FAK and AKT by transforming growth factor-β1 confers an anoikis-resistant phenotype to myofibroblasts. Cell Signal 19, 761–771 (2007).

24. Thannickal, V. J. et al. Myofibroblast differentiation by transforming growth factor-β1 is dependent on cell adhesion and integrin signaling via focal adhesion kinase. Journal of Biological Chemistry 278, 12384–12389 (2003).

25. Woodley, J. P., Lambert, D. W. & Asencio, I. O. Understanding Fibroblast Behavior in 3D Biomaterials. Tissue Eng Part B Rev 28, 569–578 (2022).

26. Cukierman, E., Pankov, R., Stevens, D. R. & Yamada, K. M. Taking Cell-Matrix Adhesions to the Third Dimension. 294, 319–342 (2012).

27. Vicens-Zygmunt, V. et al. Fibroblast viability and phenotypic changes within glycated stiffened three-dimensional collagen matrices. Respir Res 16, 1–15 (2015).

28. Karamichos, D., Lakshman, N. & Petroll, W. M. Technique article: An experimental model for assessing fibroblast migration in 3-D collagen matrices. Cell Motil Cytoskeleton 66, 1–9 (2009).

29. Uhlen, M. et al. Tissue-based map of the human proteome. Science (1979) 347, 1260419–1260419 (2015).

30. Jansen, L. E., Mccarthy, T. P. & Peyton, S. R. A synthetic, three-dimensional bone marrow hydrogel.

31. Galarza, S., Crosby, A. J., Pak, C. H. & Peyton, S. R. Control of Astrocyte Quiescence and Activation in a Synthetic Brain Hydrogel. Adv Healthc Mater 9, (2020).

32. Knight, C. G. et al. The collagen-binding A-domains of integrins alpha(1)beta(1) and alpha(2)beta(1) recognize the same specific amino acid sequence, GFOGER, in native (triple-helical) collagens. J Biol Chem 275, 35–40 (2000).

33. Patterson, J. & Hubbell, J. A. Enhanced proteolytic degradation of molecularly engineered PEG hydrogels in response to MMP-1 and MMP-2. Biomaterials 31, 7836–7845 (2010).

34. Plow, E. F., Haas, T. A., Zhang, L., Loftus, J. & Smith, J. W. Ligand binding to integrins. J Biol Chem 275, 21785–21788 (2000).

35. Lishko, V. K. et al. Multiple binding sites in fibrinogen for integrin alphaMbeta2 (Mac-1). J Biol Chem 279, 44897–44906 (2004).

36. Ruoslahti, E. RGD and other recognition sequences for integrins. Annu Rev Cell Dev Biol 12, 697–715 (1996).

37. Yamada, K. M. Adhesive recognition sequences. Journal of Biological Chemistry 266, 12809–12812 (1991).

38. Staatzs, W. D. et al. Identification of a Tetrapeptide Recognition Sequence for the alpha2 beta1 Integrin in Collagen. vol. 266 (1991).

39. Polio, S. R. et al. Cross-Platform Mechanical Characterization of Lung Tissue. PLoS One 13, (2018).

40. Barney, L. E. et al. A cell-ECM screening method to predict breast cancer metastasis. Integrative Biology (United Kingdom) 7, 198–212 (2015).

41. Reardon, D. A., Nabors, L. B., Stupp, R. & Mikkelsen, T. Cilengitide: An integrin-targeting arginine-glycine-aspartic acid peptide with promising activity for glioblastoma multiforme. Expert Opinion on Investigational Drugs vol. 17 1225–1235 Preprint at https://doi.org/10.1517/13543784.17.8.1225 (2008).

42. Hashimoto, S. et al. Transforming growth Factor-beta1 induces phenotypic modulation of human lung fibroblasts to myofibroblast through a c-Jun-NH2-terminal kinase-dependent pathway. Am J Respir Crit Care Med 163, 152–157 (2001).

43. Eickelberg, O. et al. Transforming growth factor-β1 induces interleukin-6 expression via activating protein-1 consisting of JunD homodimers in primary human lung fibroblasts. Journal of Biological Chemistry vol. 274 12933–12938 Preprint at https://doi.org/10.1074/jbc.274.18.12933 (1999).

44. Mulder, K. M. & Morris, S. L. Activation of p21(ras) by transforming growth factor β in epithelial cells. Journal of Biological Chemistry 267, 5029–5031 (1992).

45. Hanafusa, H. et al. Involvement of the p38 mitogen-activated protein kinase pathway in transforming growth factor-β-induced gene expression. Journal of Biological Chemistry 274, 27161–27167 (1999).

46. Atfi, A., Djelloul, S. H., Chastre, E., Davis, R. & Gespach, C. Evidence for a role of Rho-like GTPases and stress-activated protein kinase/c-Jun N-terminal kinase (SAPK/JNK) in transforming growth factor β-mediated signaling. Journal of Biological Chemistry 272, 1429–1432 (1997).

47. Shani, O. et al. Evolution of fibroblasts in the lung metastatic microenvironment is driven by stage-specific transcriptional plasticity. Elife 10, (2021).

48. Gocheva, V. et al. Quantitative proteomics identify Tenascin-C as a promoter of lung cancer progression and contributor to a signature prognostic of patient survival. Proceedings of the National Academy of Sciences of the United States of America vol. 114 E5625–E5634 Preprint at https://doi.org/10.1073/pnas.1707054114 (2017).

49. Scherzer, M. T. et al. Fibroblast-derived extracellular matrices: An alternative cell culture system that increases metastatic cellular properties. PLoS One 10, 1–17 (2015).

50. Oskarsson, T. et al. Breast cancer cells produce tenascin C as a metastatic niche component to colonize the lungs. Nat Med 17, 867–874 (2011).

51. Malanchi, I. et al. Interactions between cancer stem cells and their niche govern metastatic colonization. Nature 481, 85–91 (2012).

52. Talts, J. F., Wirl, G., Dictor, M., Muller, W. J. & Fässler, R. Tenascin-C modulates tumor stroma and monocyte/macrophage recruitment but not tumor growth or metastasis in a mouse strain with spontaneous mammary cancer. J Cell Sci 112, 1855–1864 (1999).

53. Saupe, F. et al. Tenascin-C Downregulates Wnt Inhibitor Dickkopf-1, Promoting Tumorigenesis in a Neuroendocrine Tumor Model. Cell Rep 5, 482–492 (2013).

54. Jensen, A. R. D. et al. Organ-Specific, Fibroblast-Derived Matrix as a Tool for Studying Breast Cancer Metastasis. Cancers (Basel) 13, (2021).

55. Gui, Y. et al. Metastatic breast carcinoma– associated fibroblasts have enhanced protumorigenic properties related to increased IGF2 expression. Clinical Cancer Research 25, 7229–7242 (2019).

56. Chung, B. et al. Human brain metastatic stroma attracts breast cancer cells via chemokines CXCL16 and CXCL12. NPJ Breast Cancer 3, 1–8 (2017).

57. Lv, Z. D. et al. Mesothelial cells differentiate into fibroblast-like cells under the scirrhous gastric cancer microenvironment and promote peritoneal carcinomatosis in vitro and in vivo. Mol Cell Biochem 377, 177–185 (2013).

58. Liu, R., Li, H., Liu, L., Yu, J. & Ren, X. Fibroblast activation protein: A potential therapeutic target in cancer. Cancer Biol Ther 13, 123–129 (2012).

59. Hocevar, B. A., Brown, T. L. & Howe, P. H. TGF-β induces fibronectin synthesis through a c-Jun pathway. EMBO Rep 18, 1345–1356 (1999).

60. Roberts, C. J. et al. Transforming growth factor β stimulates the expression of fibronectin and of both subunits of the human fibronectin receptor by cultured human lung fibroblasts. Journal of Biological Chemistry 263, 4586–4592 (1988).

61. Conte, E. et al. Effect of pirfenidone on proliferation, TGF-β-induced myofibroblast differentiation and fibrogenic activity of primary human lung fibroblasts. European Journal of Pharmaceutical Sciences 58, 13–19 (2014).

62. Cordeiro, M. F., Bhattacharya, S. S., Schultz, G. S. & Khaw, P. T. TGF-β1, -β2, and -β3 in vitro: Biphasic effects on Tenon’s fibroblast contraction, proliferation, and migration. Investigative Ophthalmology and Visual Science vol. 41 756–763 Preprint at (2000).

63. Brenmoehl, J. et al. Transforming growth factor-β1 induces intestinal myofibroblast differentiation and modulates their migration. World Journal of Gastroenterology vol. 15 1431–1442 Preprint at https://doi.org/10.3748/wjg.15.1431 (2009).

64. Pouliot, R. A. et al. Development and characterization of a naturally derived lung extracellular matrix hydrogel. J Biomed Mater Res A 104, 1922–1935 (2016).

65. Dunphy, S. E., Bratt, J. A. J., Akram, K. M., Forsyth, N. R. & el Haj, A. J. Hydrogels for lung tissue engineering: Biomechanical properties of thin collagen-elastin constructs. J Mech Behav Biomed Mater 38, 251–259 (2014).

66. Grigoryan, B. et al. Multivascular networks and functional intravascular topologies within biocompatible hydrogels. Science (1979) 364, 458–464 (2019).

67. Hynes, R. O. Integrins: bidirectional, allosteric signaling machines. Cell 110, 673–687 (2002).

68. Loffek, S., Loffek, S., Schilling, O. & Franzke, C.-W. Biological role of matrix metalloproteinases: a critical balance. European Respiratory Journal vol. 38 191–208 Preprint at https://doi.org/10.1183/09031936.00146510 (2011).

69. Engler, A. J., Sen, S., Sweeney, H. L. & Discher, D. E. Matrix Elasticity Directs Stem Cell Lineage Specification. Cell 126, 677–689 (2006).

70. Naba, A. et al. The matrisome: In silico definition and in vivo characterization by proteomics of normal and tumor extracellular matrices. Molecular and Cellular Proteomics 11, 1–18 (2012).

71. Chan, E. P., Smith, E. J., Hayward, R. C. & Crosby, J. Surface wrinkles for smart adhesion. Advanced Materials 20, 711–716 (2008).

72. Hutchens, S. B. & Crosby, A. J. Soft-solid deformation mechanics at the tip of an embedded needle. Soft Matter 10, 3679–3684 (2014).

